# Nested versus independent sampling: Solving the mystery of contradictory species-area relationships

**DOI:** 10.1101/2020.08.19.257337

**Authors:** Johannes Reinhard, Barbara Drossel

## Abstract

Species-area relationships (SARs) describe how the number of species increases with the size of the area surveyed, and they usually take the shape of a power law on regional spatial scales. A meta-review of empirical data has shown that the exponent of the power law is on average larger when the areas are sampled in a nested manner, compared to sampling of independent areas such as islands of different sizes. As this is in contrast to ecological reasoning, we performed computer simulations of three qualitatively different models that generate species distributions in space and time by the mechanisms of speciation, dispersal, and extinction. We find that in all cases and over a wide parameter range the SARs obtained by nested sampling have a smaller slope in the regional scale than those obtained by independent sampling. We explain the discrepancy to the empirical data by the different spatial scales on which the two types of empirical investigations were performed.

## 1 Introduction

Ecological systems are the result of a long-term evolutionary process that involves speciation, dispersal, and extinction, leading to ever changing local, regional, and global species compositions. The exploration of the species composition on different spatial scales has revealed a variety of patterns that appear to be the generic outcome of this long-term process. These patterns include rank-abundance distributions, species-area relationships, and distance-similarity relations. Among these, species-area relationships (SARs) are probably the ones that have been investigated for the longest time, as they involve merely counting the number of species *n_S_* of the class of interest found within areas of different sizes *A*. The data reveal a monotonically increasing function the shape of which was proposed by Arrhenius [1] to be a power law

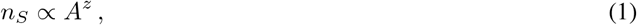

while Gleason [2] proposed a semi-logarithmic relation *n_S_* = *a* + *b* log*A*. Of these two laws, the power law is more widely accepted in the literature [3], and it is often a better fit to the data, as reviewed by Drakare et al. [4] Of course this law is not valid over all scales but only on intermediate scales. As discussed by Rosenzweig [5] and Hubbell [6], the SAR is a triphasic species-area curve, summarized by Drakare et al. [4] as follows: ‘At local scales (within communities), SARs are driven by rank-abundance distributions and are nonlinear in log–log space. At intermediate scales, SARs depend less on abundance patterns and more on speciation, dispersal and extinction, and give a linear relationship in log–log space. At even larger geographical scales (crossing evolutionary provinces), assemblages do not share evolutionary history and are not connected by dispersal, leading to an increase in steepness of slopes as species-assemblages are unrelated and species turnover maximized.’ This triphasic curve is is shown schematicallyin fig. 1(a). A power law of the form (1) occurs only in the intermediate range, where the slope is approximately constant in this log-log plot. Fig. 1(b) shows directly the slope, which decreases at the local scale, is constant on the regional scale and increases towards 1 on the continental scale.

**Figure 1:**
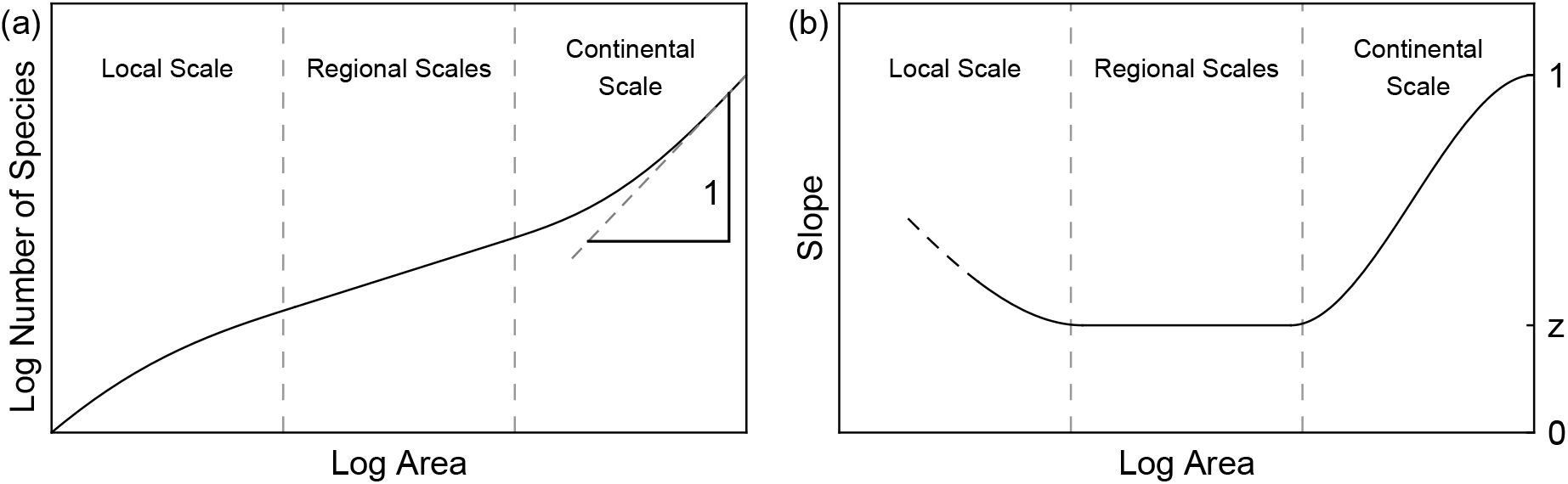
Schematic diagram of a) the triphasic species-area curve in a log-log plot and b) the slope of the curve shown in (a). Diagram (a) is drawn according to Hubbell [6].

In the last 100 years there have been numerous empiric studies that either investigated SARs directly or that collected data that are useful for evaluating SARs. There a several different ways for evaluating how the number of species increases with the considered area [7], with the two most important ones being nested and independent sampling. In case of nested sampling, the areas of increasing sizes *A*_1_, *A*_2_,… are chosen such that the area with the next size *A_n_* fully contains the previous area of size *A*_*n*−1_. In the case of independent sampling, the areas of different sizes are independent from each other and (usually) not overlapping, such as islands or different nature reserves.

Drakare et al. [4] collected the remarkable amount of 794 of these studies in a meta-analysis. This analysis includes a variety of aquatic and terrestrial ecosystem types. Taking all the studies together, they obtained an average *z*-value of 0.27. When evaluating separately the slopes obtained with nested and independent sampling methods, they found that the mean *z*-value for nested sampling was 0.36, while it was 0.24 for independent sampling. This finding is surprising as one would intuitively expect the opposite behavior, as mentioned for instance by Rosenzweig [5, 8] and illustrated by fig. 2, which shows the two possible relationships between nested and independent SARs. If we assume that nested sampling is done within each island used for independent sampling, the end point of each nested SAR must be coincide with the data point for independent sampling for this island size. Logically, there are two possibilities: Either the nested SARs are above the independent SAR, or they are below the independent SAR. In the first case, one expects the nested SAR exponent to be on average smaller than the independent SAR exponent, in the second case one expects the reverse. Now, we would expect that larger islands host within the same area more species than smaller islands. This means that the nested SARs should lie above the independent SARs, as represented in fig. 2(b), and in contradiction to fig. 2(a) and the findings by Drakare et al. [4]

**Figure 2:**
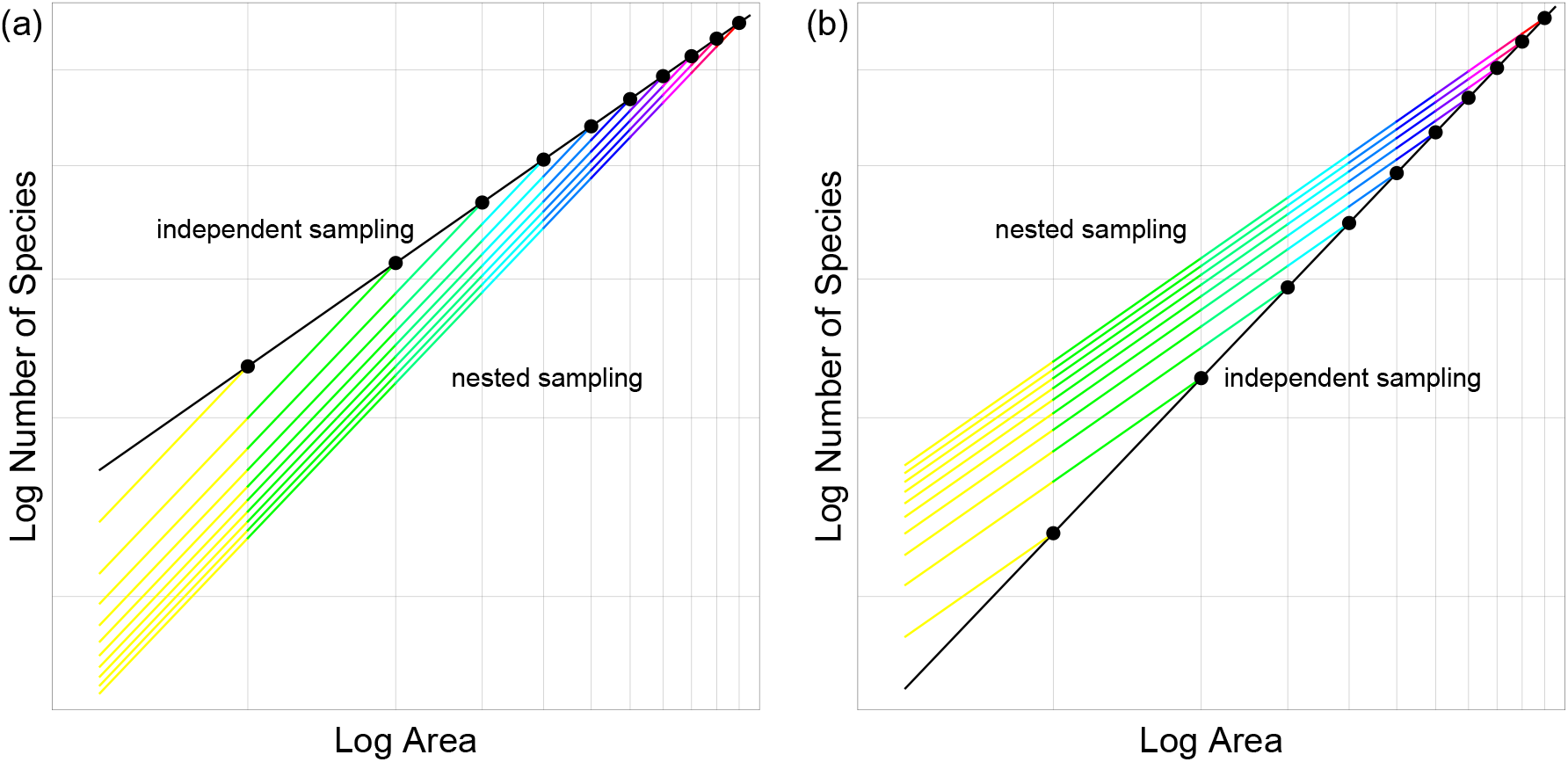
The two possible qualitative relationships between the SARs obtained from nested and independent sampling. (a) The nested sampling curves lie below those for independent sampling and are on average steeper. (b) The nested sampling curves lie above those for independent sampling and are on average flatter.

It is the purpose of this paper to resolve this puzzle by analysing several theoretical models that are based on the combined processes of speciation, dispersal, and extinction and generate species distributions in space. We will use three different models that differ with respect to the spatial scales and the complexity of the ecosystems. The first model is the neutral, individual-based model by Hubbell [6] where each site is occupied by one individual. The second model is the evolutionary meta-food web model by Rogge et al. [9] that resolves space only on the level of habitats and leads to the formation of complex trophic networks in each habitat. The third one is a hybrid model that considers the basal trophic layer on the level of habitats, each of which can host a given number of species.

We will evaluate all these models by comparing the results of independent and nested sampling. In fact, various authors have investigated SARs and other ecological laws by performing computer simulations of such models [9, 10, 11], but none of them has evaluated the SAR by independent sampling or compared both sampling methods.

Computer simulations of all three models will confirm the scenario depicted in fig. 2(b), i.e., the SARs obtained from independent sampling are below those obtained from nested sampling. But we will also see that the slopes of the independent sampling SARs change considerably with spatial scale and do not show the triphasic behavior of fig. 1. Together with the change in slope of the nested sampling SARs, we will find that on smaller spatial scales that do not yet represent the regional scale the slope of the nested sampling SAR may lie above that of the independent sampling SAR.

Finally, we will show that the resolution of the puzzle posed by the findings by Drakare et al can be found by considering the spatial scales on which the different empirical studies were performed. Since the nested sampling studies were on average performed on much smaller spatial scales than the independent sampling studies, the SAR exponents *z* were larger than they would have been if one had evaluated them on the scale on which the independent sampling studies were performed.

## 2 Models and methods

### 2.1 The individual-based neutral model (Hubbell model)

The first model is a spatially explicit version of Hubbell’s neutral model [6] as used by Rosindell and Cornell [10], but with a fixed system size. The model uses a square lattice of *n* nodes called sites, each of which is inhabited by one individual. Every individual belongs to a species, which is labeled by an integer index.

At each step of the simulation, a site *m* is chosen at random and the individual on this site is removed. Then one of two possible events occurs, weighted by their rates:

- **Speciation event** with rate *μ_s_* = 1: A new species is introduced into the system, and an individual of this species is placed on lattice site *m*.
- **Dispersal event** with rate *μ_d_*: One site *l* is chosen at random from all sites within a square dispersal kernel of width 2*L* +1 around site m, and a copy of the individual on site *l* is placed on site *m*. This individual might belong to the same species as the removed individual.

This model is best interpreted as describing the trees in a forest. About a quarter of the studies from Drakare et al. [4] focus on the ecosystem type ‘forest’, which deals mostly with basal species, in particular with trees. Therefore, the Hubbell model is particularly suitable as a computer model for this class of empirical data.

### 2.2 The evolutionary model

The second model is the spatially explicit evolutionary meta-food web model introduced by Rogge et al. [9] It generates meta-ecosystems with local food webs in each habitat, and it is therefore appropriate for computer simulations of SARs for systems with several trophic levels.

The model is defined on a square lattice with n nodes, which now represent habitats, each containing its own local food web and an external resource *R*.

Each species *S* is characterized by three traits, which determine its feeding links. These traits are the body mass *m_S_*, feeding center *f_S_* and feeding range *s_S_*. A species *S* can feed on species *P* (or the external resource *R*) if log *m_P_* (or log *m_R_* ≡ 0) is within the interval log *f_S_* ± *s_S_*. The local instance of a species in a habitat is called a population of this species. A population *i* can be present or absent but has no actual population size. It can survive only as long as its survival index

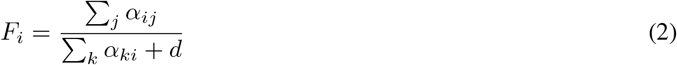

is larger than or equal to 1. The survival index depends on the link strength *α_ij_* to predator population *j* and on a loss term *d*. The link strength is given by

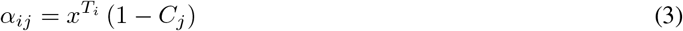

where *x* < 1 models efficiency losses, *T_i_* is the trophic level, and

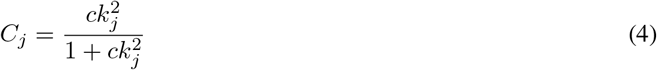

reduces the link strength due to competition between the *k_j_* predators of population *j*. The trophic level of a population *i* is recursively defined by the average trophic level of the prey of population *i* plus 1, with the trophic level of the resource being zero by definition.

The simulation is started with the resource on each habitat and with one species that has traits such that it can feed on the resource. In each step of the simulation, one population *i* is chosen at random out of all populations on all habitats, and one of the following three events occurs with a probability weighted by its rate:

- **Speciation event** with rate *μ_s_* = 1: A new Species *C* is created that is a modification of the parent species *P* to which the chosen population belongs. Its body mass and feeding center are chosen at random from the intervals *m_C_* ∈ [0.2 *m_P_*, 5 *m_P_*] and *f_C_* ∈ [0.001*m_C_*, 0.1*m_C_*], and its feeding range is fixed to *s_C_* = 0.5. A population of this species is added to the habitat of the parent population, and its survival index (2) is evaluated. If it is below 1, the speciation event was not successful and the new population is removed from the food web. Otherwise, the new survival index of all other populations is evaluated. If the survival index of one ore more populations is smaller than 1, the population with the smallest survival index goes extinct and is removed from the food web. (if several populations have the same survival index, one is chosen at random.) This procedure is repeated until no survival index is below 1.
- **Dispersal event** with rate *μ_d_*: A habitat within a square dispersal kernel of width 2*L* +1 around the habitat of population *i* is chosen randomly. A new population of species *P* is inserted into this habitat. If there is already a population of this species or if its survival index is less than 1, the dispersal event was not successful and the new population is removed from the food web. Otherwise the survival indices of all other species are evaluated. If at least one of them is below 1, the extinction process described above is repeated until the survival indices of all remaining populations are ≥ 1.
- **Spontaneous extinction event** with rate *μ_e_*: Besides extinction events triggered by the addition of new populations, there are also spontaneous extinction events, which occur with a small rate. In this case, the chosen population is removed from the habitat, possibly leading to the extinction of further populations according to the process described above. This spontaneous extinction rate is required to prevent a freezing of the system.

Computer simulations of this model lead to meta-food webs with several trophic layers, which show an ongoing change of the species composition and their distribution over the habitats. After an initial phase where the food webs increase in size, a stationary state is reached where the population numbers fluctuate around some mean value. In this stationary state, SARs can be evaluated.

### 2.3 The hybrid model

The third model is a hybrid of the first two models. As in the Hubbell model, no traits are assigned to species apart from their index, as they are all equivalent. In contrast to the Hubbell model, the nodes of the lattice in the hybrid model are habitats instead of sites for only one individual. Just as in the evolutionary model, populations are not assigned a population size, but are simply present or absent. Each habitat can host *ω* distinct populations. In the limit case *ω* = 1, this model becomes almost identical to the Hubbell model, except for minor details in the dispersal process. For *ω* > 1, its SARs differ qualitatively from the Hubbell model, as we will see in the Results section. This model can be interpreted as an extension of the Hubbell model to much larger spatial scales and to patchy landscapes, where distinct habitats are connected by dispersal. During each step of the simulation, a population *i* out of all populations on all habitats is chosen at random, and one of the following two events occurs with a probability weighted by its rate:

- **Speciation event** with rate *μ_s_* = 1: A new species is created, and its first population replaces population *i*.
- **Dispersal event** with rate *μ_d_*: A random population *j* is chosen from the habitats within a square dispersal kernel of width 2*L* + 1 around the habitat of population *i*, and a copy of it is placed into that habitat instead of population *i*. If the species to which population *j* belongs is already present in that habitat, the dispersal event is not successful.

### 2.4 Parameter values used in the simulations

The three models share several parameters, but there are also parameters that occur only in one of the models. The parameter values respectivly the ranges within which they were varied are listed in Tab. 1.

**Table 1:**
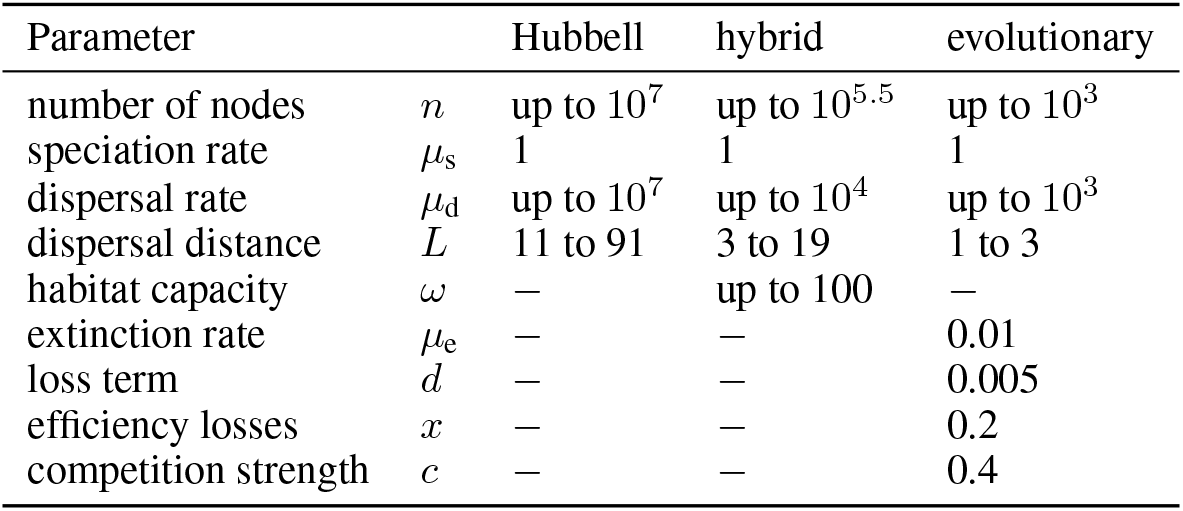
Parameters of the models

| Parameter |  | Hubbell | hybrid | evolutionary |
| --- | --- | --- | --- | --- |
| number of nodes | $n$ | up to $10^7$ | up to $10^{5.5}$ | up to $10^3$ |
| speciation rate | $\mu_s$ | 1 | 1 | 1 |
| dispersal rate | $\mu_d$ | up to $10^7$ | up to $10^4$ | up to $10^3$ |
| dispersal distance | $L$ | 11 to 91 | 3 to 19 | 1 to 3 |
| habitat capacity | $\omega$ | — | up to 100 | — |
| extinction rate | $\mu_e$ | — | — | 0.01 |
| loss term | $d$ | — | — | 0.005 |
| efficiency losses | $x$ | — | — | 0.2 |
| competition strength | $c$ | — | — | 0.4 |

The parameters *x*, *c*, and *d* of the evolutionary model were chosen according to the standard parameter set of Rogge et al. [9], which gives food webs of a feasible size. The dispersal rate *μ_d_*, the dispersal distance *L*, the system size *n*, and the habitat capacity *ω* of the hybrid model were varied in the indicated ranges.

The dispersal rate *μ_d_* = 10^5^ corresponds to that with which most of the results of Rosindell and Cornell [10] were generated. The dispersal distances *L* = 11 and 91 were also examined there.

### 2.5 Simulation procedure

We performed computer simulations of all three models using the programming language C++. For the Hubbell model, we implemented the algorithm by Rosindell et al. [12], which runs the dynamics backwards in order to construct one system snapshot that is part of the stationary state. For each parameter set, the number of different snapshots was at least as large as 100000 divided by the number of patches.

For the hybrid model and the evolutionary model, we performed 10 long simulation runs for each parameter set, which allowed us to evaluate the species-area relationship at several different points in time. In order to obtain good statistics, a simulation duration of 1 mutation event per habitat and local population, and an evaluation of the SAR at five points in time was sufficient for the hybrid model, while the simulation time was 50000 speciation events per habitat for the evolutionary model (with most of these not leading to the survival of the new species), evaluating the SAR in intervals of 200 speciation events per habitat. Since the evolutionary model requires considerable time until the stationary food-web size is reached, we started evaluations only after 8000 speciation events.

For all models, nested sampling was performed starting from the central patch and adding in each step as many patches as are required to obtain a square of the next size, centered around the starting patch.

In order to plot the slope of the SARs, the difference quotient of log(*S*) and log(*A*) was evaluated.

## 3 Results

Fig. 3 shows the species-area relationships and the corresponding slopes for all three models for one choice of the parameters each. For the nested SAR curves, the original data points (small, non-black dots) were logarithmically binned to obtain the solid lines. This procedure is also used in all further figures, and the individual data points will not be shown any more.

**Figure 3:**
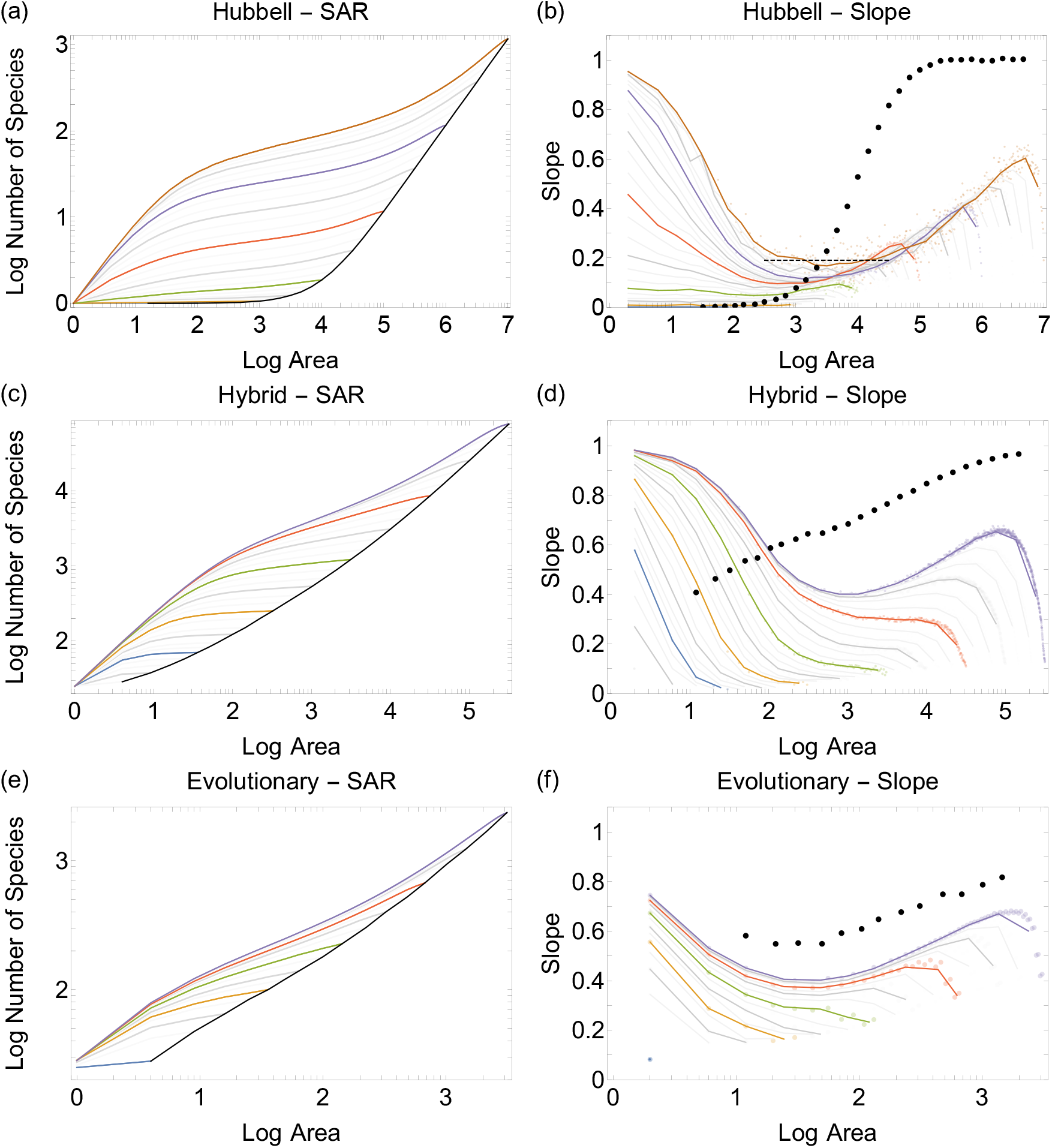
SARs and corresponding slopes for the Hubbell model ((a),(b); *μ_d_* = 10^5^, *L* = 11), the hybrid model ((c),(d); *μ_d_* = 10^3^, *L* =11, *ω* = 25, and the evolutionary model ((e),(f); *μ_d_* = 10^2^, *L* = 2). Colored and gray curves refer to nested sampling, with the color allowing easy identification of a pair. The black curves in (a), (c), (e) refer to independent sampling and connect the end points of the nested sampling SARs. The black dots in (b), (d), (f) give the corresponding slopes.

The first information to be obtained from this figure is that the nested SAR curves are always above the independent SAR curves, thus confirming the scenario depicted in fig. 2(b). This is also the case for all later simulations, for which we will only show the slopes as they are sufficient to convey all remaining information.

Second, the local, regional and global scales cannot be as clearly distinguished as fig. 1 suggests. There is no section where the slope really remains constant. The regional scale is in the range around the minimum of the slope but has no clear boundaries.

Third, with increasing system size (which can be read off from the end points of the different nested SAR curves) the curves approach an asymptotic curve for area values sufficiently far below the total area. For an infinite area, Rosindell and Cornell [10] gave the value 0.19, which is indicated by the dashed horizontal line in fig. 3(b) and is approached by the curves for the larger system sizes.

Fourth, the independent SAR curves for the Hubbell and the hybrid model do not show the expected triphasic behavior, with the difference to this behavior being most strongly pronounced in the Hubbell model.

In the following we investigate how the dispersal rate and the dispersal distance affect these results. For the hybrid model, we explored also the effect of the habitat capacity. However, as all values *ω* > 1 gave almost identical results, we do not show these data. As already mentioned, the data for *ω* = 1 are very similar to those of the Hubbell model and need therefore not be shown either.

Fig. 4 shows the influence of dispersal rate and dispersal distance on the SARs for all three models. For each model, a low and a high dispersal rate and a smaller and larger dispersal distance were chosen. Each of the figures shows the slope of the independent SAR curve and the largest nested SAR curve examined.

**Figure 4:**
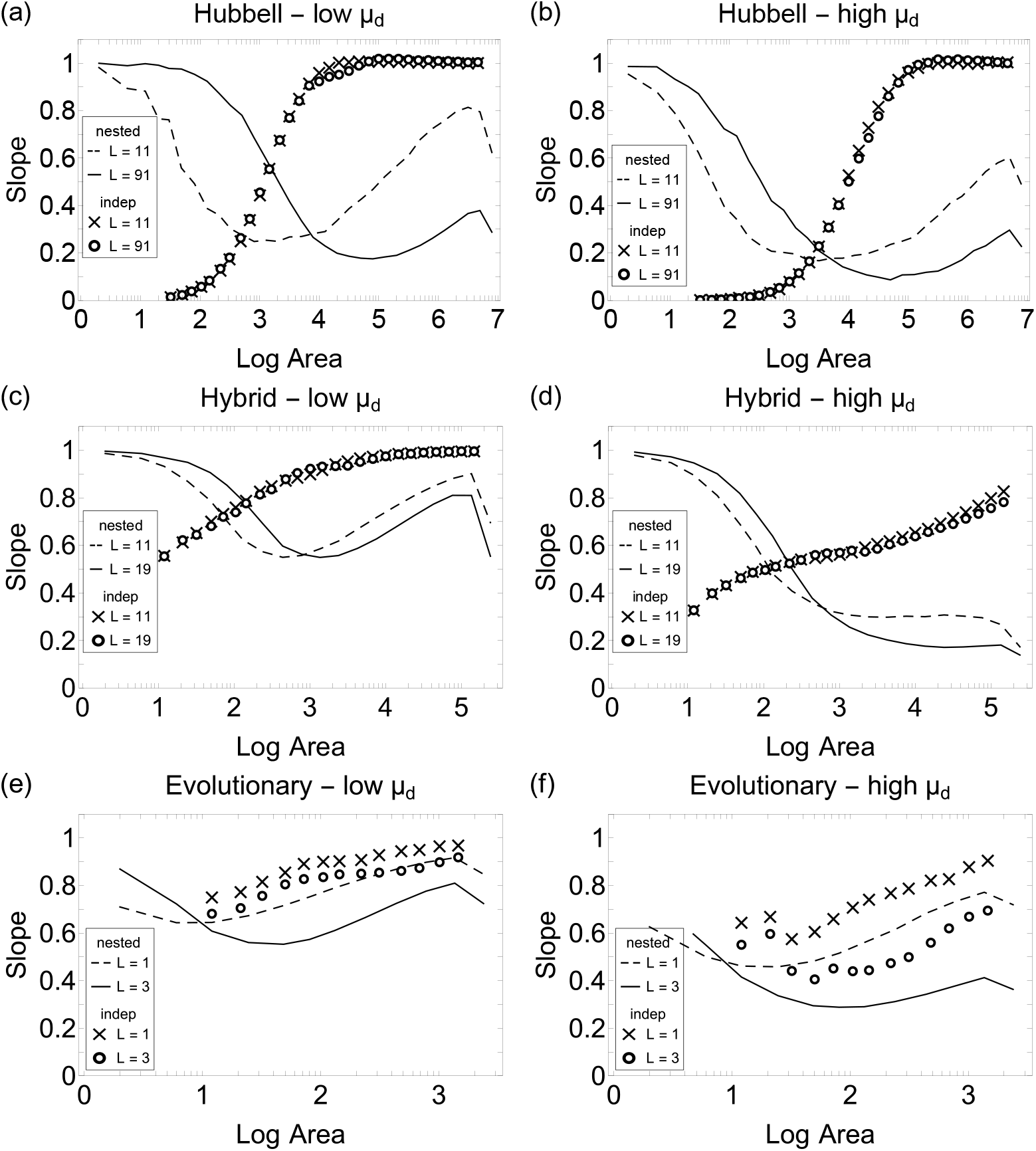
The slope of the SAR for the Hubbell model ((a),(b); *μ_d_* = 10^5^, *L* = 11, 91), the hybrid model ((c),(d); *μ_d_* = 10^3^, 10^4^, *L* = 11, 19, *ω* = 25, and the evolutionary model ((e),(f); *μ_d_* = 10^1^, 10^3^, *L* = 1, 3). Curves refer to nested sampling, crosses and circles refer to independent sampling For the nested SARs, only the curves for the largest system sizes are given.

Let us first consider the nested SARs. They show the expected triphasic shape described in fig. 1, apart from the hybrid model with large dispersal rate, where finite-size effects are apparently so strong that the final increase is not visible. In an infinite system, all slopes must eventually reach the value 1.

The main effect of dispersal distance *L* and dispersal rate *μ_d_* on the nested SARs is that they move to the right when one of these parameters increases. This can be understood from the fact that species that have an *n*-fold dispersal distance or dispersal rate can reach an *n*^2^-fold area in the same time. This shift has also been described by Rosindell and Cornell [10] for the Hubbell model. The fact that the slopes for large area decrease with increasing dispersal distance and dispersal rate is due to finite-size effect, as the cutoff in system size has a stronger effect when dispersal distances are larger.

The behavior of the independent SARs is more complex and differs between the three models. The most striking feature is that the dispersal range does not affect the SAR for the two neutral models. This is explained as follows: The number of species in a system depends only on how often new species enter the system, which results directly from the ratio of dispersal rate and speciation rate. The dispersal distance only determines the course of the nested SAR up to the maximum species number and this is irrelevant for the independent SAR.

This does not apply to the evolutionary model as this is not a neutral model. Extinction and dispersal processes depend on the spatially varying food web structure, and therefore these two processes do not occur completely randomly as in the neutral models. In this situation, a larger dispersal distance conveys higher survival chances, and therefore the independent SARs move to the right with increasing dispersal distance in the evolutionary model.

The shape of the independent SARs differs between the three models. For the Hubbell model, only a single species exists in the system for small system sizes and large dispersal rates. In the range of these system sizes the slope is zero. Additional species that are introduced by speciation processes can survive only for a short time, leading to a very slow initial increase of the slope with increasing system size. On the other end of the curves, the system size has become so large that the number of species is proportional to the system size, leading to a slope of 1. In between these two limits there is a transition zone that moves to larger areas with increasing dispersal rate.

The hybrid and the evolutionary model do not show the initial stage of slope 0 of the independent SARs. This is because each habitat now hosts several species, and it becomes very unlikely that all habitats contain the same set of species. For the hybrid model, this is because a species can only be driven out of a habitat by another species if this other species is not already present the habitat. In the evolutionary model, the different structure of different local food webs hampers the global spreading of species. These two models would therefore need much larger dispersal rates in order to reach the situation that all habitats host the same set of species.

Finally, and most importantly, all three models show that in general the slopes of the independent SARs lie above those of the nested SARs, except for small spatial scales that are not yet in the regional range. Only for the Hubbell model in the case of large dispersal rates and small dispersal distances do the slopes of the nested and independent SAR curves intersect in the regional scale.

During the course of this research, we have also investigated other variants of these systems, such as Random Geometric Graphs instead of regular lattices, or periodic instead of open boundary conditions, but this did not lead to qualitatively different results or to new insights. The use of a different dispersal kernel had already been investigated by Rosindell and Cornell [10] for the Hubbell model, finding that this has no qualitative influence on SARs.

## 4 Discussion

Our investigation of three conceptionally different models and a wide range of parameter values has unambiguously shown that the nested SARs lie always above the independent SARs, and that the slopes of the nested SARs on regional spatial scales are smaller than the independent SARs in almost all cases. Only for the Hubbell model and only for the unrealistic combination of very large dispersal rates and small dispersal distances do the slopes of the independent and nested SAR curves intersect on the regional scale.

As this is in contrast to the empirical data reviewed by Drakare et al. [4], we will in this concluding section discuss the reasons for this discrepancy. To this purpose, we have taken a closer look at the data used by Drakare et al. [4] and examined the sizes of the areas surveyed. The result is shown in Figure 5, once for all ecosystem types (a), and once only for the ecosystem type ‘forest’ (b). Each study is represented by a bar, which extends from the smallest investigated area to the largest investigated area. When nested SARs were examined in a study, the bar is red; when independent SARs were considered, the bar is blue. In addition, the area averaged over all studies of a survey method is shown as a dashed vertical line of the respective colour. The heights of the horizontal dashed lines indicate the respective average slopes, showing the values of z given by Drakare et al. [4]. From these plots, is evident that the independent and nested studies were in general conducted on different scales. When averaging over the studies for all ecosystem types, nested SAR data were gathered on areas of approximately 20m^2^ and independent SAR data on 330, 000m^2^. For the ecosystem type ‘forest’, these two areas are 170m^2^ and 580, 000m^2^. There are therefore 4.2 or 3.5 orders of magnitude in between the two types of SAR data. Furthermore, various sources [5, 13, 14, 15] suggest that the regional scale for many species starts at the earliest in the 10^4^m^2^ range. Specifically for trees, the curves of Plotkin et al. [16] show that before 10^5^m^2^ the regional scale is not yet reached. Only the fewest studies with nested SARs surveyed by Drakare et al. are thus within the regional scale. Mostly, these data were obtained on the local scale. But on the local scale, slopes are significantly steeper than on the regional scale, cf. fig. 1 and our simulation results in figs. 3 and 4. We illustrated this in fig. 5(c), where we represented the data from fig. 3(d) in the same way as the data reviewed by Drakare et al. [4] This picture shows that even in our computer simulations, the nested data for small areas are above the independent ones for larger areas, even though we know that the nested corves lie above the independent curves, and that there slopes are smaller than those of the independent SARs when both are taken on the regional scale.

**Figure 5:**
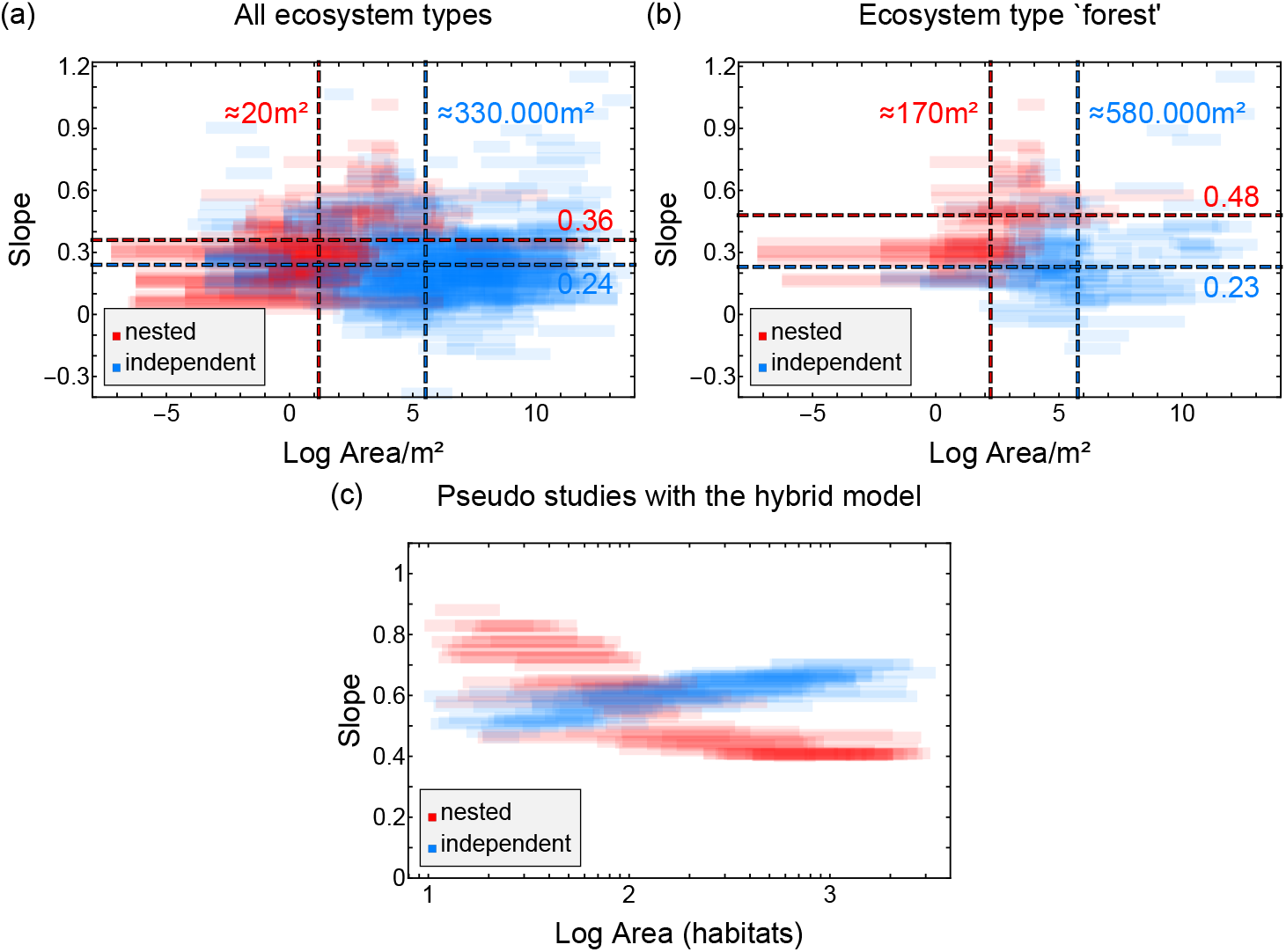
Surveyed areas and the corresponding slopes of the studies used by Drakare et al. [4] (a) all studies (b) studies of ecosystem type ‘forest’ (b). In (c) we show data generated from fig. 3(d) by selecting random intervals for areas of “pseudo studies”.

All this means that the discrepancy between theory and expectation on one hand and empirical data on the other hand is due to the limitations of the available empirical data. It is not easy to make a study large enough to count species numbers on nested areas of square kilometers and larger. Moreover, nested and independent sampling have never been carried out on the same system, so that the two types of studies are difficult to compare. In our computer simulations, we could overcome both these limitations, as we could perform both types of studies on the same spatial scales and by using exactly the same type of ‘ecosystem’. Our work thus has demonstrated the usefulness of theoretical models for gaining a deeper understanding of empirical data and for finding explanations for puzzling observations.

## Acknowledgements

We thank Kay-Robert Dormann for extracting the data for Fig 5 from the literature and the DFG for funding via the project Dr300/12-2 and Dr300/13-2.

